# CRISPR-Cas9 editing of grapevine clade V-related *mlo* genes revealed new insights into the contribution of each *mlo* gene to grapevine powdery mildew resistance (*Erysiphe necator*) and plant development

**DOI:** 10.64898/2026.01.29.702653

**Authors:** Satyanarayana Gouthu, Laurent G. Deluc

## Abstract

MLO proteins are a family of transmembrane proteins that play an essential role in plant immunity, host-pathogen interactions, and a wide range of other developmental processes in land plants. More specifically, clade V-MLOs are associated with powdery mildew susceptibility in several plants. In grapevine, four MLO genes, *MLO3*, *4*, *13*, and *17,* belong to clade V and have been studied to determine their contribution to the susceptibility to *Erysiphe necator* (Grape Powdery Mildew - GPM). Using somatic embryogenesis to generate knockout mutants in grapevine, a series of multiplex gene-editing-mediated transformations was performed to determine the contribution of each clade V-related MLO to GPM resistance. A hierarchy of contributions from the four MLOs to GPM was characterized, with *MLO17* having the most significant role, followed by *MLOs 3* and *13*. In addition, *MLO4* was shown to be a non-essential modifier of GPM resistance: its loss-of-function, in combination with other MLOs, conferred a modest but consistent improvement in GPM resistance. However, defects in flower development in the majority of mlo4 mutant lines suggest a role for this gene in reproductive tissues. Our observation also indicated functional redundancy within the clade V-MLO, particularly between MLO13 and MLO17. Ultimately, our results show that a certain degree of chimerism (wild-type background in the mutants) is necessary to achieve relatively complete resistance to GPM without major growth defects or yield penalties.

## Introduction

MLO (mildew Locus O) family proteins have been identified in several plant species, including 17 MLOs in grapevine (Feechan et al., 2008), and some of them have been shown to impact infection by fungi causing powdery mildew (PM) (Consonni et al., 2006a; Dong-Yan et al., 2020; Ingvardsen et al., 2019; Pessina, Lenzi, et al., 2016a). The MLO orthologs responsible for PM susceptibility are conserved across plant species and generally cluster in clades IV and V (Feechan et al., 2008; Kusch & Panstruga, 2017; Pozharskiy et al., 2022). MLOs are required for pathogen-host cell penetration and negatively regulate plant defenses against fungal invasion. MLO proteins colocalize with and directly or indirectly influence the plasma membrane-localized soluble N-ethylmaleimide sensitive factor attachment receptor (SNARE) penetration-resistance proteins (AtPEN1, AtPEN3, HvROR2, VvPEN1), which deposit cell wall reinforcements to act as physical barriers against fungal penetration (Bhat et al., 2005; Feechan et al., 2013; Underwood & Somerville, 2008). Loss of function at the MLO locus confers resistance to the powdery mildew pathogen by obstructing the pathogen’s entry (Underwood & Somerville, 2008).

*mlo*-based resistance against powdery mildew is naturally found or can be introduced by silencing MLO genes in a broad range of plant species (Acevedo-Garcia et al., 2017; Consonni et al., 2006a; Pessina, Angeli, et al., 2016; Zheng et al., 2013). Beyond modulating PM resistance, MLO family proteins are involved in a wide range of physiological processes, including calcium signaling, root morphogenesis, and responses to biotic and abiotic stresses (Bidzinski et al., 2014; Chen et al., 2006; Kim et al., 2002). As a result, both natural and induced loss-of-function mutations in *MLO* genes produce pleiotropic effects, such as spontaneous necrotic leaf spotting, increased susceptibility to other diseases (McGrann et al., 2014), and reduced yield, among others. Therefore, despite being attractive targets for plant breeding programs to introduce PM resistance, the use of *mlo*-based resistance in agriculture is limited primarily due to the yield penalty from growth defects associated with mutations. However, two studies have demonstrated that the severity of growth defects can be reduced by partially restricting MLO expression and by residual MLO wild-type activity, the natural variant of mlo11 in barley (Ge et al., 2020; Piffanelli et al., 2004). Likewise, gene-edited chimeric mutant of *edr1* (Enhanced resistance1), another PM susceptibility gene in grapevine, expressing wild-type and mutant alleles, was reported to develop normally because of lower editing frequency in meristem tissues compared to the older tissues that display PM resistance (Yu et al., 2024).

In grapevine, *MLO* genes involved in PM susceptibility have been investigated based on their homology to functionally validated mildew susceptibility genes *AtMLOs 2, 6, and 12* (Consonni et al., 2006b), which in turn, are co-orthologs of the natural barley *mlo* allele (Panstruga, 2005b). In *Arabidopsis,* complete protection against PM requires the silencing of all three clade V MLOs, 2, 6, and 12, due to high functional redundancy. Early grapevine studies focused on the identification of *MLO* genes involved in PM infection through their induction kinetics (Feechan et al., 2008; Winterhagen et al., 2008), followed by loss-of-function or knockdown studies (Dong-Yan et al., 2020; Moffa et al., 2024; Pessina, Lenzi, et al., 2016b). Complementation experiments with grapevine MLOs 3 and 4 partially rescued PM susceptibility phenotype in PM-resistant *Arabidopsis* triple mutant, *Atmlo2,6,12,* and both were also shown to co-accumulate at the site of fungal penetration, along with PEN1 (Feechan et al., 2013). Grapevine RNAi experiments suggested a 77% reduction in PM severity associated with knockdown of *MLOs 3*, *13*, and *17* and no role for *MLO4* (Pessina, Lenzi, et al., 2016b). Using gene-editing approaches, the *VvMLO3* mutation has been reported to confer partial resistance to PM, accompanied by growth defects (Dong-Yan et al., 2020). Near-complete resistance was reported with double mutations in *MLOs 13* and *17,* without pleiotropic effects, attributed to overaccumulation of stilbenes and improved oxidative scavenging (Moffa et al., 2024).

In this study, gene-edited lines carrying combinations of clade V *MLO* mutations were generated to identify PM-susceptibility candidates and to assess the potential combinatorial effects on PM resistance. Based on resistance tests, mutant genotyping, and phenotype observations, the relationship between *MLO* gene combinations and resistance to powdery mildew, as well as the associated developmental defects caused by the loss of MLOs is partially elucidated. The results suggest that unintended chimerism, often associated with somatic embryogenesis following gene editing, might have alleviated the pleiotropic effects while retaining the benefits of *mlo*-based resistance in grapevine.

## Materials and Methods

### Microvine plant material and embryogenic callus

The microvine 04C023V0004 genotype used in this study, progeny from a cross between ‘Grenache’ × L1 ‘Pinot Meunier’ GAI1 mutant, described elsewhere (Chaïb et al., 2010), was provided by Dr. Mark Thomas (CSIRO Laboratories, Australia). The microvine plants were grown in a greenhouse under 16-h day and 8-h night conditions provided by supplemental lighting (100 μM m-2s-1, equivalent to 5,400 lux). Initiation of embryogenic callus from immature anthers, maintenance media, and culture conditions were performed as described previously (Torregrosa et al., 2015).

### Guide RNAs and design of CRISPR-Cas9 constructs

CRISPR-Cas9 constructs were designed to generate single knockouts (*mlo 3*, *mlo 4*, *mlo 13*, and *mlo 17)*, double knockouts (*mlo3,4* and *mlo13,17),* and a quadruple knockout (*mlo3,4,13,17*) using the CRISTA algorithm to rank sgRNA (Abadi et al., 2017). The last CRISPR construct was also intended to generate any incidental combinations of double (*mlo 4,13* and *mlo 4,17*), and triple mutant combinations (*mlo3,4,13*, *mlo3,4,17*, *mlo4,13,17*, *mlo3,13,17*). Unique single guide RNAs (sgRNA) for single knockout mutants of *MLO*s 3, 4, 13, and 17 were selected from the 2^nd^ or 3^rd^ exon regions. A common sgRNA was designed for *MLO*13 and *MLO17* for the double knockout *mlo13,17*, whereas individual sgRNAs of *MLO3* and *MLO4* were cloned into the CRISPR vector for the *mlo3,4* double mutant. Plasmid for *mlo3,4,13,17* quadruple mutant (*qko*) was designed by multiplex cloning the sgRNAs of MLOs 3 and 4, and the common sgRNA of MLOs 13 and 17 into the plasmid (**Fig. S1**). The single-transcriptional-unit (STU) CRISPR/Cas9 vector, pGEL031 (Addgene Plasmid #137900), was used with the eGFP gene to screen for transformed embryos. In the STU vector, Cas9 and sgRNAs are expressed from the maize ubiquitin Pol II promoter, and multiple sgRNAs are spaced with tRNAs to separate the individual sgRNAs post-transcription (Tang et al., 2019).

For single knockouts, sgRNAs were synthesized as duplex oligonucleotides, annealed, and cloned into BsaI-linearized pGEL031. To create double knockouts, the sgRNA scaffold was amplified by PCR using a pair of specific primers including two sgRNA sequences (BsaI-sgRNA1-F and BsaI-sgRNA2-R), and the purified PCR product was cloned. To express three sgRNAs for quadruple knockout, a two-sgRNA fragment and a third common sgRNA generated through PCR amplification, along with the tRNA spacer sequence, were together cloned into pGEL031 through Gibson assembly.

### Microvine Transformation

*Agrobacterium*-mediated transformation was performed following established protocols for the microvine embryogenic callus (Torregrosa et al., 2015). Briefly, embryogenic calli of microvine 04C023V0004 were co-incubated with *Agrobacterium tumefacien*s EH105 harboring the pGEL031 vector containing each sgRNA or combinations of sgRNAs (**Table S1**), and the infected calli were maintained on embryo-inducing medium with hygromycin selection. Six to eight weeks after the transformation, the emerging somatic embryos were screened for GFP fluorescence, and the GFP-positive embryos were regenerated. Regenerated transformants were propagated and maintained in magenta boxes for genotype confirmation and further analyses.

### Genotyping and Mutation Analysis

For mutant genotyping, the genomic DNA was extracted from the leaf tissue of regenerated plants using DNeasy Plant Mini kit (Qiagen). PCR reactions were prepared using 50 ng genomic DNA as the template and specific primers designed to amplify a 200-300 bp amplicon spanning the expected double-strand break site of the target MLO gene. PCR products were purified using the Qiagen PCR purification kit (Qiagen) and sequenced using a nested primer (**Table S2**). Online analysis of mutant genotypes was performed by submitting the ab1 files from Sanger sequencing to the DECODR deconvolution tool to detect indels (Bloh et al., 2021). The PCR amplicons were sequenced using Oxford Nanopore long-read sequencing by Plasmidsaurus (Eugene, OR, USA). Approximately 5,000 reads per sample were obtained from NGS and were analyzed using the CRISPR edits analysis tool (Geneious Prime, 2023.2.1).

### Powdery mildew strain and leaf inoculation

*Erisyphe necator*, isolate E1-101 was maintained on detached leaves of Cabernet Sauvignon. To inoculate microvine leaves for PM resistance studies, conidia from 8-day-old colonies were used. Leaves of tissue-culture-grown microvine plants were transferred to water agar medium in Petri dishes, with their petioles inserted into the medium and a piece of wax paper placed over the leaves to prevent direct contact with the agar. For the induction experiment, conidia were deposited on the entire leaf surface using a small paintbrush (size 5/0) and incubated at 25 °C until the specified collection times.

### MLO-induction time-course studies

To test for induction of MLO genes upon PM inoculation, the leaves were uniformly coated with conidia using a paintbrush. RNA was extracted from the inoculated leaves, and cDNA was synthesized and used as a template for real-time PCR. Relative expression of MLOs was calculated using the ΔΔCt method (Livak & Schmittgen, 2001).

### Mycelial growth and sporulation area analysis

For the resistance studies, conidia were deposited on each side of the midvein and between the first and second lateral veins of a detached leaf (as above). Stereoscopic observations of initial spore germination and hyphal emergence were done 24 hpi. Mycelial growth and sporulation were scored at 72 hpi. Traces of the leaf area, the total area of the leaf with mycelial spread, and the area under sporulation were made on the lids of Petri dishes. The images were uploaded to ImageJ, and the fungal growth (cm^2^) was calculated as a percentage of total leaf area (Abràmoff et al., 2004).

### MLO-induction time-course studies

To test for induction of MLO genes upon PM inoculation, the leaves were uniformly coated with conidia using a paintbrush. RNA was extracted from the inoculated leaves, followed by cDNA synthesis to was used as a template for real-time PCR. Relative expression of MLOs was calculated using the ΔΔCt method (Livak & Schmittgen, 2001).

### Real-time PCR for quantitative fungal biomass analysis

Assessment of fungal biomass on infected leaves was performed 2 weeks post-inoculation (wpi) in a separate experiment using leaves spot-inoculated with *E. necator* conidia as described above. Quantitative assessment of fungal biomass in infected leaves was performed using real-time PCR to determine the ratio of fungal to microvine genomic DNA (Webling & Panstruga, 2012). Genomic DNA was extracted from the infected leaves using the DNeasy Plant Mini Kit (Qiagen) and used as the template for real-time PCR with primers for the *E. necator* 18S ribosomal RNA gene (GeneBank: LC777882) and the *Vitis* actin gene. For qPCR, 20 μl reactions were prepared using the SsoAdvanced Universal SYBR Green Supermix Kit (Bio-Rad) according to the manufacturer’s protocol. The amplification of the *E. necator*-derived amplicon was normalized to the microvine actin gene amplicon to quantify fungal biomass. The ratio of *E. necator* to microvine genomic DNA in mutants was expressed relative to the WT ratio.

### Real-time PCR experiments for MLO homologue expressions

To test for compensatory expression of MLO homologs in mutants, the leaves were spot inoculated with *E. necator* conidia, and the inoculated leaves were collected at 72 h and 2 wk post-inoculation at specified times. RNA was extracted from the samples, and cDNA was synthesized for use as a template in real-time PCR. Relative expression of MLOs was calculated using the ΔΔCt method (Livak & Schmittgen, 2001). The primers used in real-time PCR are listed in Table S1.

### Statistical analyses

qRT-PCR expression data of the four MLO genes in control and infected leaves were analyzed using a standard least-squares linear model with genotype and infection status as fixed effects. Relative expression was calculated by the ΔΔCt method and expressed as fold change relative to non-infected wild-type samples and to the *MLO3* gene at 72 hpi in control and infected leaves. Mutant genotypes were compared with the wild type and the *MLO3* gene using least-squares means with Dunnett’s multiple-comparison test (α = 0.05). Data represent means ± SEM of five biological replicates. Fungal growth and sporulation were quantified at 72 hpi by measuring the proportion of leaf area covered by *Erysiphe necator* mycelium and sporulating structures. Measurements were obtained from 10–14 independently infected leaves per genotype. Because the data did not meet the assumptions of normality and homoscedasticity, differences between *mlo* mutants and wild-type plants were evaluated using pairwise nonparametric Wilcoxon rank-sum tests. A similar design was performed for the biomass estimate analyses. For the 72 hpi gene redundancy experiment, relative gene expression was calculated using the ΔΔCt method and expressed as fold changes relative to non-infected wild-type (WT) samples. For each gene, expression data were analyzed using a linear model including genotype and infection status as fixed effects. Least-squares means were estimated, and mutant genotypes were compared to WT using Dunnett’s multiple comparison test. Differences between non-infected and infected samples within each genotype were assessed using model-based contrasts. Data represent means of five biological replicates ± SEM, and statistical significance was evaluated at α = 0.05. For the 2 wpi gene redundancy experiment, data were analyzed for each gene and response variable, using a linear model including *mutant* as the main fixed effect and *date* as a blocking factor to account for batch effects (*y* ∼ Clone + Date). Model inference was performed using heteroskedasticity-consistent covariance estimators (HC3) to ensure robustness to non-constant variance. Model assumptions were evaluated using diagnostic plots. Pairwise comparisons among clones were conducted using Tukey’s post-hoc tests based on robust standard errors, and results were summarized using compact letter displays.

## Results

### Clade V mlos in grapevine and induction during powdery mildew infection

Using phylogenetic analysis, four grapevine MLOs were identified as the closest orthologs of functionally characterized PM-susceptibility MLOs from *Arabidopsis* and other species (Feechan et al., 2008, Winterhaggen et al., 2008). Two different gene annotations are currently used by the scientific community from the papers cited above (**Table S3**), we opted for Feechan’s annotation in this manuscript, *i.e.* MLO 3, 4, 13, and 17. These four MLOs share the two conserved short C-terminal domains common to all MLOs, with confirmed roles in powdery mildew susceptibility in other dicot species (Panstruga, 2005a; Humphry et al., 2011; Kim and Hwang, 2012; Yan et al., 2021). Our microvine genome shows that *MLO13* and MLO*17* are adjacent to each other, suggesting their paralogous nature (Unpublished data) and that they have highly similar protein sequences.

Transcript levels of the four candidate MLOs were and quantified to determine how their early- and late-stage responses to PM inoculation with established infection stages of *Erysiphe necator* (**Table S4**). The induction patterns of *VvMLOs 3*, *4*, *13*, and *17* in infected microvine leaves were observed 4 hours post-inoculation (hpi) to 7 days post-inoculation (dpi) (**Fig. 1a**). While the early-stage transcriptional responses varied among the four MLO homologs, they were generally more expressed during colony expansion (3-7 dpi). At 2-4 hpi period (conidial germination and attempted leaf penetration), *MLOs 4* and *17* gene expression showed a significant ∼4-to-5-fold increase (**Fig. 1a**). At 24 hpi, when spores from haustoria and primary hyphae have formed, no significant changes were observed in the expression of the four *mlo* genes. At 3 dpi, when conidiophore development begins, only *MLO13* was highly expressed. At 7 dpi, *MLO4* and *MLO17* again showed significantly higher expression during this stage (**Fig. 1a**). *MLO3* expression was largely unaffected by the infection process, or its induction was not captured during our sampling times. Given the steady-state pattern of *MLO3*, we use it to evaluate the relative expression levels of *MLOs* in healthy and infected leaves at 3 dpi (**Fig. 1b**). In non-infected leaves, *MLO4* expression was at a similar level to that of *MLO3,* but *MLOs 13* and *17* were expressed at a 4-fold higher rate. Relative expression rate for *MLOs 13* and *17* was even higher in infected leaves at 40 to 50-fold compared to *MLO3* (**Fig. 1b**). Previous tissue-specific expression studies suggest that *MLOs 3* and *4* are predominantly expressed in floral and root tissues, indicating their role in developmental functions beyond PM interaction (Feechan et al., 2008; Winterhagen et al., 2008), and the pleiotropic effects we observed in the mutants, discussed later in the manuscript, support this notion.

**Figure 1:**
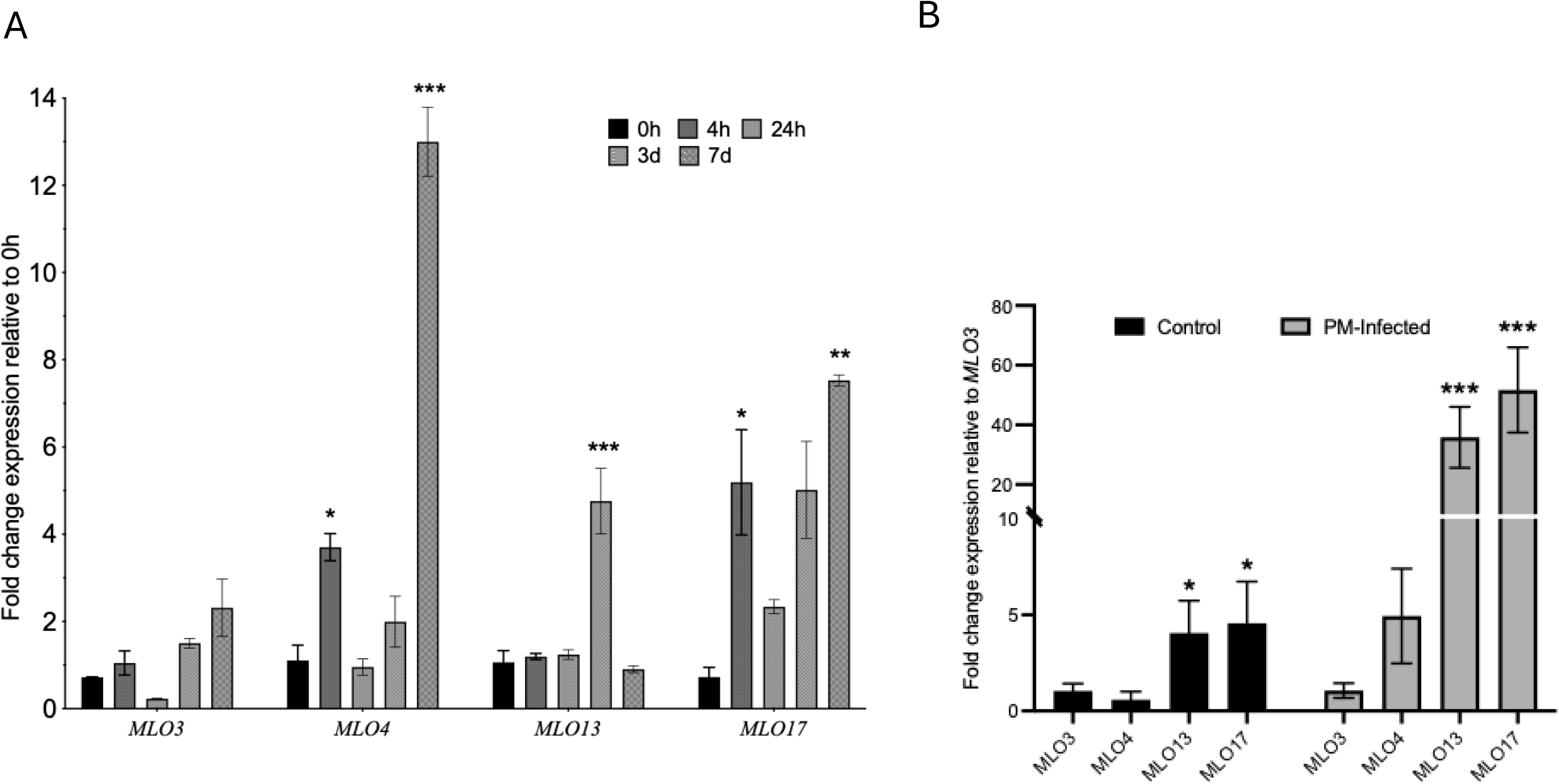
Real-Time PCR analysis of *VvMLO* gene expression in grape leaves in response to powdery mildew infection. **A.** Expression fold changes were calculated relative to the 0h sample for each *VvMLO* gene. Means were calculated from 3 to 7 biological replicates. Error bars show SEM, and an asterisk indicates a statistically significant difference from their respective 0h samples as determined by Dunnett’s test. 0h (un-inoculated) detached leaves were incubated the same way as with 4h inoculated leaves. **B.** Expression fold changes in control and infected leaves at 72hpi relative to *VvMLO3* gene expression. Means were calculated from 3 to 7 biological replicates. Error bars show SEM, and an asterisk indicates a statistically significant difference from *VvMLO3* expression values as determined by Dunnett’s test. *: p<0.05, **: p<0.01, ***: p<0.001

### CRISPR-Cas9 editing performance, genotyping, wild-type chimerism, and pleiotropic effects

PM susceptibility is attributed to the overlapping roles of clade V MLOs such as AtMLOs 2, 6, and 12 in *Arabidopsis* (Consonni et al., 2006a) and SlMLOs 1, 5, and 8 in tomato (Yan et al., 2021). To identify the contribution of each MLO to PM susceptibility, CRISPR-Cas9 constructs were designed to generate single-, double-, triple-, and quadruple-knockout lines (**Fig. S1 and Table S1**). *In vitro* cleavage assays confirmed the specificity of each sgRNA-Cas9 construct with no off-target cleavage among the other MLO homologs (**Fig. S2**). Following the transformation, we observed embryo clusters during embryogenesis with fused basal parts, which were avoided whenever isolated embryos were available. Five non-retrieved knockouts (*mlo4,13*, *mlo4,17*, *mlo3,4,13*, mlo3,4,17, *mlo4,13,17*) had an sgRNA targeting MLO4, which showed a decent editing rate in the single knockout construct but failed to perform in the other CRISPR constructs. The last non-retrieved knockout was *mlo13,17*, which could also be explained by an overall lower *in planta* editing performance of the sgRNA targeting MLO17 (**Table S6**). We also incidentally retrieved double and triple mutants *mlo3,13* and *mlo3,13,17* while screening for quadruple knockout lines. The transgenic lines were evaluated for mutations mainly using the deconvolution tool DECODR (Bloh et al., 2021) (**Table 1**). Of the 15 potential *mlo*-knockout combinations that could be generated from our CRISPR construct designs, we identified 9 (**Table 1**). The amplicon sequencing performed on *mlo3*, *mlo4*, *mlo13*, *mlo17*, and *qko #1* lines was consistent with DECODR analyses within the 5% error margin (**Fig. S4**) and validated the presence of mosaicism editing and WT chimerism in the mutants. Bi- or monoallelic mutations were determined by the complete absence of the wild-type (WT) allele or ∼50% contribution of the WT allele. We classified the mutation types as homozygous, heterozygous, or mosaic editing (more than two edited allele types) (**Table 1-Table S5**). Mutants in which the WT MLO allele contribution was too far from the halfway mark are classified as chimeric plants. Overall, *mlo3* and *mlo4* were biallelic and completely edited lines, whereas *mlo13* and *mlo17* were chimeric, with genotype showing ∼75% of the reads contributed by the unedited WT MLO allele (**Table 1**). Among double mutants, *mlo13,17* was chimeric for *MLO17*. In double mutants of *mlo3,4* and *mlo3,13,* the *MLO3* mutation was biallelic, whereas its counterparts were monoallelic. Both quadruple mutants, *qko #1* and *#2*, were chimeric for *MLO4* and *MLO17*, respectively (**Table 1**, **Fig. 2**). Among the incidental knockouts (*mlo3, 13* and *mlo3, 13, 17*), both showed mono-, biallelic, and mosaic editing (**Table 1**, **Fig.2**). The numbers of confirmed edited lines out of the potential transformants show that sgRNAs of *MLO3* and *MLO13* had higher editing efficiency (Fig. S5) compared to MlO4 and MlO17, which explained the absence of mutant combinations with *MLOs 4* and *17*.

**Figure 2:**
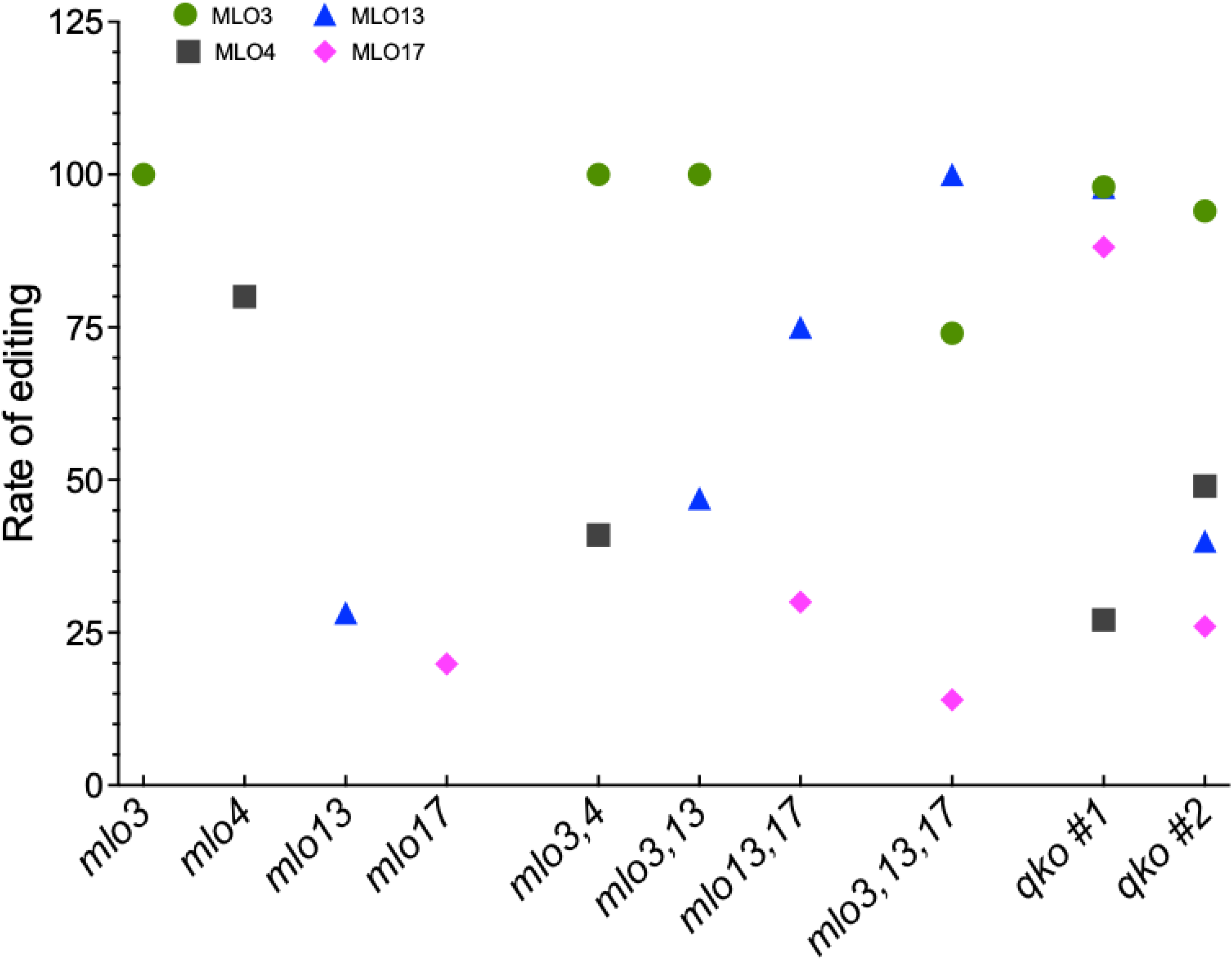
Rate of editing in the major mutants identified in this study. The editing mutation rate was calculated from the percentage of mutated alleles detected in the genotype analysis relative to the Wild-Type sequence, as determined using DECODR (Bloh et al., 2021).

**Table 1:** Characterization of *mlo* mutant genotypes (DECODR and Long-Read sequencing) along with their observed phenotypic pleiotropic effects. The mutants were categorized based on the type of editing on the alleles, the type of resulting mutation, and the presence of WT-related chimerism. Type of editing: Mono/Biallelic refers to one or both alleles edited. Mutation type: Homozygous refers to the same kind of mutation in both alleles, while heterozygous refers to different types of mutations in alleles; Mosaic refers to more than two types of editing; Chimeric refers to the presence of unedited wildtype sequences along with edited sequences. –: refers to a level of editing that was below a biallelic or monoallelic expected percentage.

| Mutant | Target gene | Type of editing | Mutation type | Plant | Phenotype in soil conditions |
| --- | --- | --- | --- | --- | --- |
| *mlo3* | 3 | Bi | Homozygous |  |  |
| *mlo4* | 4 | Bi | Homozygous |  | Slow growth, Non-flowering |
| *mlo13* | 13 | - | Mosaic | Chimeric with WT | Severe leaf necrotic lesions upon soil transfer, Normal development after juvenile stage |
| *mlo17* | 17 | - | Mosaic | Chimeric with WT | Stunted growth. Normal development after juvenile stage |
| *mlo3,4* | 3 | Bi | Homozygous |  | Slow growth, Leaf senescence, Non-flowering |
|  | 4 | Mono | Heterozygous |  |  |
| *mlo3,13* | 3 | Bi | Homozygous |  |  |
|  | 13 | Mono | Mosaic |  |  |
| *mlo13,17* | 13 | Mono | Mosaic |  | Black roots, loss of root gravitropism, narrow and thick leaves. Didn't survive past 4<sup>th</sup> micropropagation. |
|  | 17 | - | Mosaic | Chimeric with WT |  |
| *mlo3,13,17* | 3 | Bi | Heterozygous |  | Slow growth and nonflowering |
|  | 13 | Bi | Mosaic |  |  |
|  | 17 | Mono | Mosaic | Chimeric with WT |  |
| *qko #1* | 3 | Bi | Mosaic |  |  |
|  | 4 | - | Mosaic | Chimeric with WT |  |
|  | 13 | Bi | Mosaic |  |  |
|  | 17 | Bi | Mosaic |  |  |
| *qko #2* | 3 | Bi | Homozygous |  | Stunted growth, premature senescence, did not survive |
|  | 4 | Mono | Heterozygous |  |  |
|  | 13 | Mono | Heterozygous |  |  |
|  | 17 | - | Mosaic | Chimeric with WT |  |

In terms of pleiotropic effects, most mutants exhibited slow growth to severe growth defects, including arrested growth and non-flowering phenotypes, resulting in the loss of mutants during tissue culture or under greenhouse conditions. *mlo13,17* showed major defects in root growth and couldn’t survive under greenhouse conditions (**Table 1**). *qko#2* was severely stunted, with premature leaf senescence in tissue culture, and did not survive the transfer to greenhouse conditions (**Table 1**). Only three mutant lines, *mlo3*, *mlo3,13,* and *qko#1,* lacked pleiotropic effects and showed growth, flowering, and fruit development identical to those of WT plants (**Table 1**).

### Powdery mildew resistance in mlo mutants

Resistance to PM was assessed based on mycelial growth and sporulation, and fungal biomass in infected leaves (**Figs. 3, 4 and S3**). Early microscopic observations of WT leaves inoculated with *E. necator* show that the majority of conidia had germinated and produced primary hyphae by 24 hpi (**Fig. S3**). By 72 hpi, mycelial growth covered about 20-45% of leaf area, with 40-90% of it producing conidiophores (**Fig. 3**). *mlo3* and *mlo4* showed similar levels of conidial germination and fungal growth, but the sporulation in *mlo3* was at a reduced level compared to WT (**Fig. 3**). In *mlo13* and *mlo17* mutants, hypervirulent hyphal growth was observed with significantly larger leaf area covered by powdery mildew compared to WT (**Fig.3**). The rapid conidial germination and fungal growth in these mutants were evident with secondary hyphae production at 24 hpi (**Fig. S3**). *mlo13* and *mlo17* mutants are partially edited WT chimeric lines in which the majority of cells contain WT *MLO* alleles (**Fig. 2**), and it is unclear how partially edited *MLOs* cause hypersensitivity to PM. Double mutation in *MLOs 3* and *4* caused increased hyphal spread area in *mlo3,4* but the mycelial growth was less dense with a significantly reduced sporulation (**Fig.3**). Mutation in *MLOs 3* and *13* resulted in decreased hyphal growth and unaffected sporulation in *mlo3,13* (**Fig. 3**). By contrast, *mlo13,17* and *mlo3,13,17* showed increased mycelial spread in infected leaves compared to WT. The quadruple mutant line, *qko #2*, with partially edited *MLO17*, supported WT levels of fungal growth but significantly inhibited sporulation, similar to *mlo3,4*. Finally, the quadruple mutant line, *qko #1*, with completely edited *MLOs 3, 13,* and *17* for partially edited *MLO4*, showed complete inhibition of fungal growth (**Fig. 3**). No conidial germination and emerging primary hyphae were seen in this mutant at 24 hpi (**Fig. S3**). Out of multiple qualitative resistance trials with *qko #1*, only two isolated small-colony appearances were observed, confined to the leaf base and not spreading further. In conclusion, inhibition of fungal sporulation appears to be associated with mutations in *MLO3* and *MLO4*, but an additional *MLO17* mutation also reduces hyphal growth.

**Figure 3:**
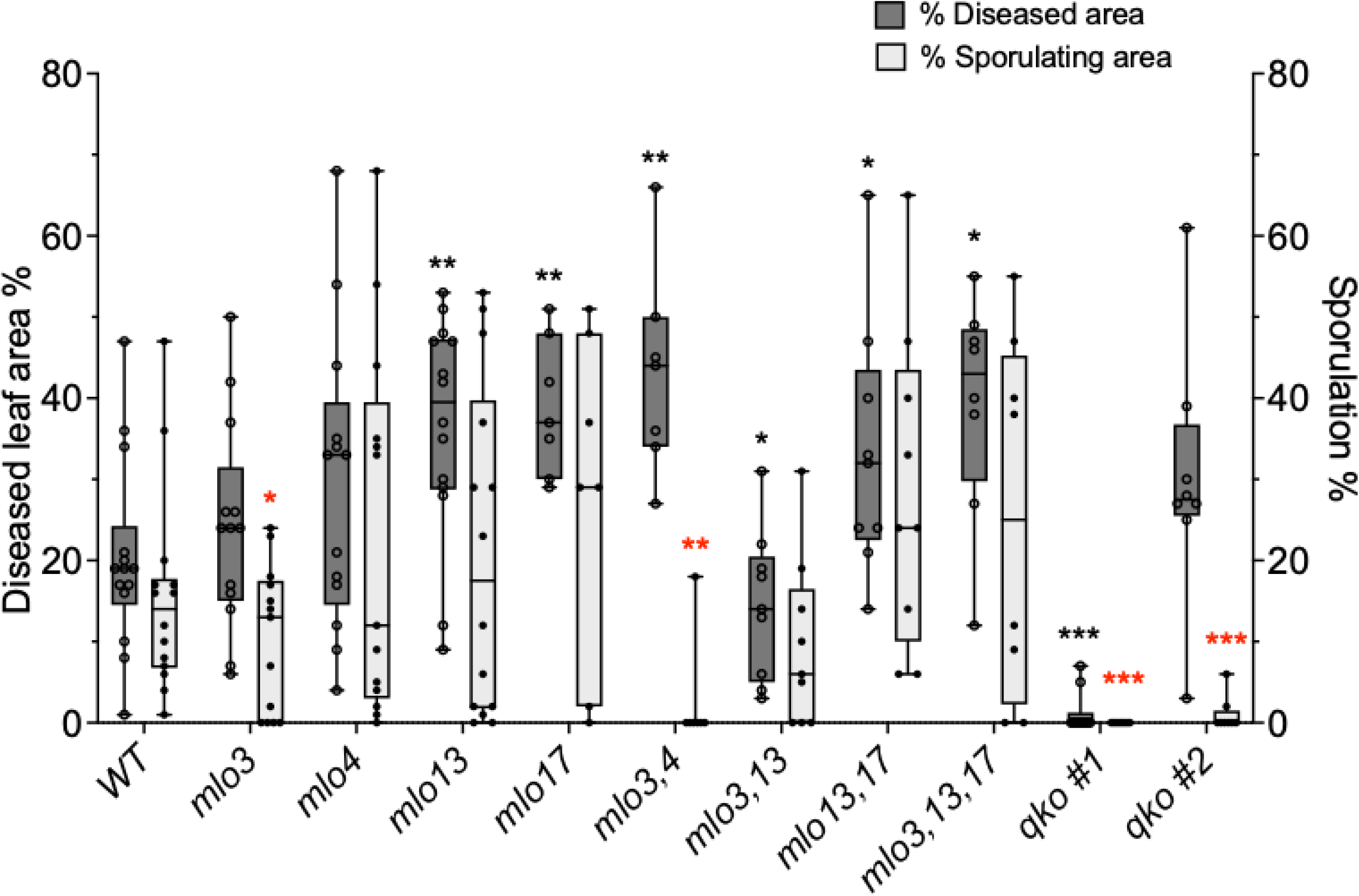
Proportion of the *E. necator* hyphal growth and sporulation areas in the *mlo* mutants. Area (cm^2^) of the total leaf area covered by mycelial growth and the sporulating area of fungal growth were acquired using ImageJ software by making the traces of the leaf outline, the areas covered by mycelial growth, and conidiophore growth 72 hours after inoculation with *E. necator* conidia. Data from 10 to 14 infected leaves was shown. Results of pairwise nonparametric comparisons using the Wilcoxon rank-sum test were presented with black and red asterisks indicating significant differences in fungal growth per leaf area and percentage sporulating area, respectively, in *mlo* mutants compared to WT leaves. *: p<0.05, **: p<0.01, ***: p<0.001

To quantitatively determine the fungal biomass supported by infected leaves at 14 dpi, we used the ratio of *E. necator* genome-derived UNC gene and the grapevine actin gene. Realtime PCR data indicate no reduction in fungal biomass in any of the single mutants compared to WT leaves. Consistent with observed hypervirulent growth, *mlo13* and *mlo17* showed a significant increase in fungal biomass (**Fig. 4**). In general, mutations in multiple MLO homologs (double and triple KO) resulted in lower fungal biomass with significant reduction for *mlo3,4*, *mlo3,13,* and *qko#2*. The quadruple mutant line *qko#1*, with partially edited *MLO4,* supported negligible fungal biomass. These results indicate that achieving complete PM resistance may require mutations in all four MLOs in a specific ratio, with MLOs 3, 13, and 17 playing a significant role.

**Figure 4:**
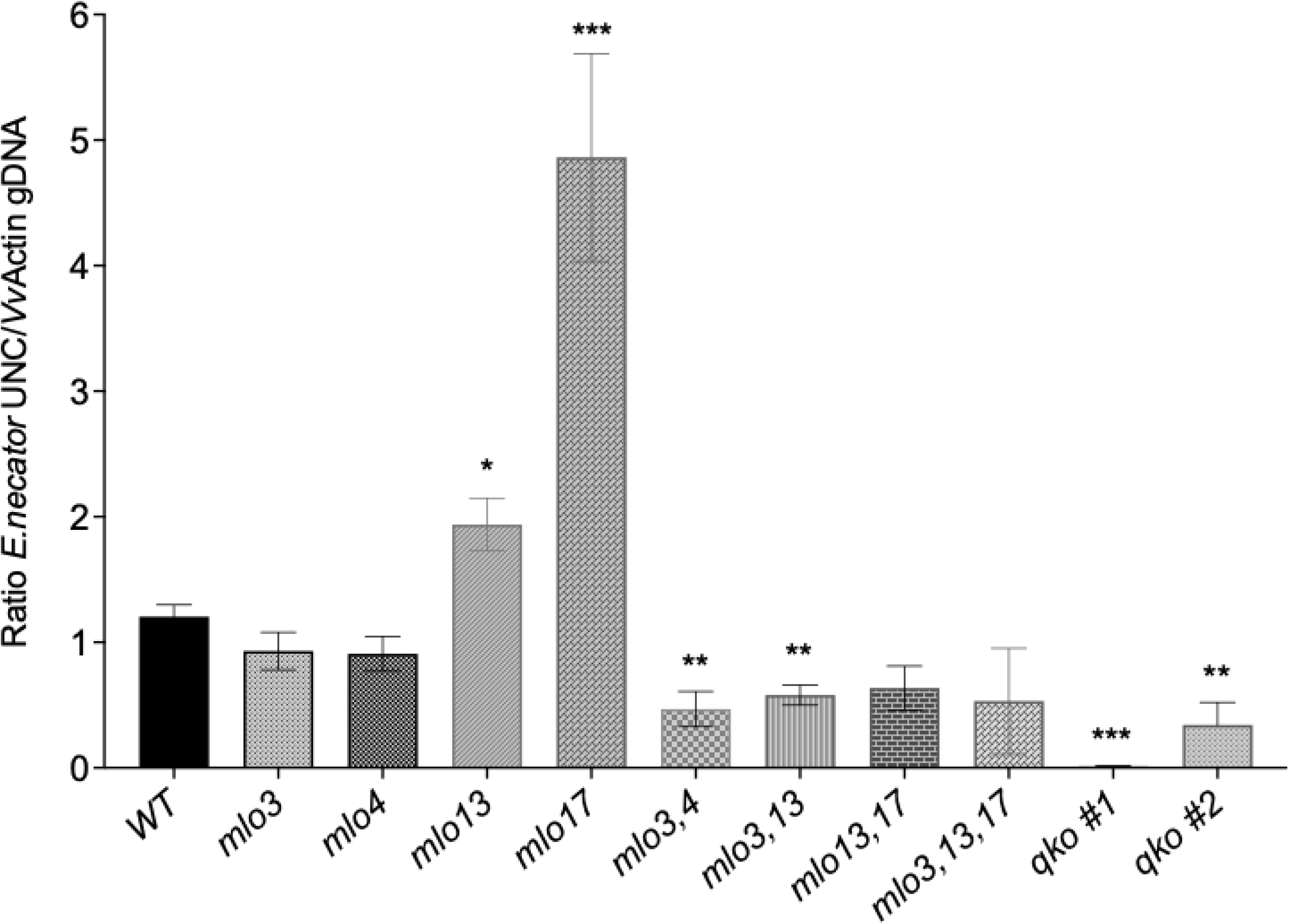
Evaluation of Erysiphe necator biomass in mutants after two weeks post inoculation (wpi). Ratios of *E. necator* to microvine gDNA in the leaves were determined quantitatively by qPCR with *E. necator* UNC gene and *Vitis* Actin gene primers using gDNA of infected leaves 2 wpi. Data represent the average of 7-12 leaves collected from 4 to 5 plants per experiment, across three independent inoculation experiments. Error bars represent SEM. Significant differences in fungal biomass in *mlo* mutants compared to WT were calculated through pairwise nonparametric comparisons using the Wilcoxon rank-sum test and were shown with asterisks. *: p<0.05, **: p<0.01, ***: p<0.001.

### Expression patterns of MLO orthologs in the mutants

Transcriptional responses of the other *MLO* homologs in single knockout and the *qko#1* mutant line were examined to determine any compensatory expression mechanism among the homologs as a result of known function redundancy among MLOs during PM infection (Consonni et al., 2006b, 2010). Early responses were examined at 72 hpi in non-infected and infected mutants and WT (**Fig. 5**). Long-term transcriptional responses were measured 2wpi (**Fig. 6**). No significant expression changes in MLO genes were observed in mutant lines compared to WT in the absence of PM infection, indicating that the loss-of-function in one of the MLOs did not cause any compensatory response in other clade v homologs (**Fig. 5 A-D** grey bars). When challenged with PM, the expression of *MLO3* did not change significantly in response to the loss of its homologs in *mlo4*, *mlo13*, and *mlo17* lines (**Fig. 5A**). *MLO4* did not express in response to PM challenge in WT but was expressed at significantly higher levels in *mlo3* and *mlo13* in response to the loss of corresponding MLOs (**Fig. 5B**). PM infection induced a higher expression of *MLO13* in WT, but the expression increase was at a much more pronounced level in *mlo3*, *mlo13*, and *mlo17* mutants (**Fig. 5C**), indicating that MLO13 might compensate for the loss-of-function in MLOs 3 and 17. Under pathogen-challenged conditions, *MLO17* was induced in WT, but in all the mutants except *mlo3, MLO17* expression was at significantly lower levels (**Fig. 5D**). While the loss of MLOs 3, 13, and 17 in single mutants seemed to be compensated by one or more homologs, it was not apparent in the quadruple mutant, *qko#1,* besides substantially lowered expression of *MLO13* and *MLO17* compared with WT under pathogen-challenged conditions.

**Figure 5:**
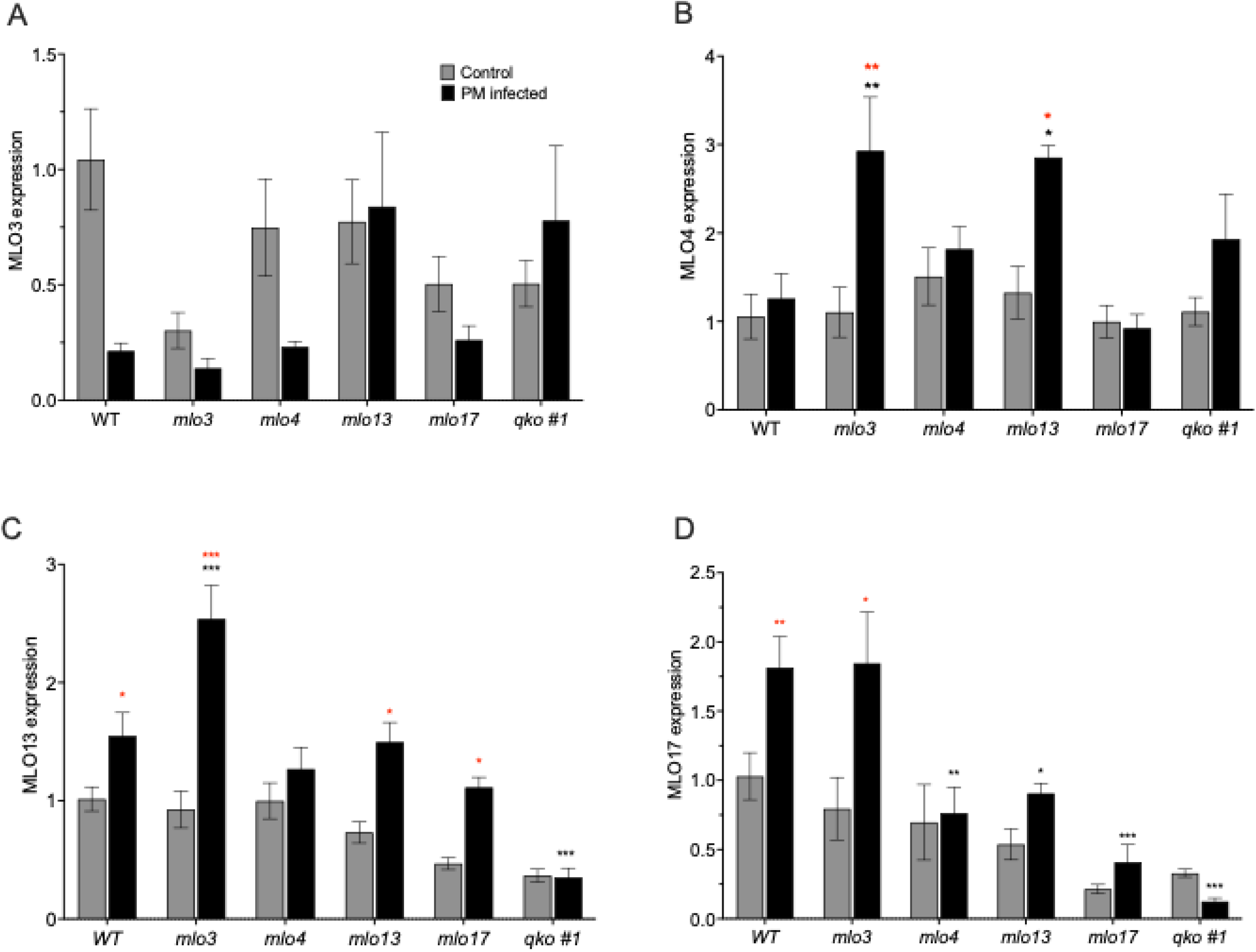
Early transcriptional response of MLO orthologs in non-infected and powdery mildew-infected *mlo* mutants 72 hpi quantified through quantitative RT-PCR analyses. The results are expressed as fold changes relative to the expression in non-infected WT. The black asterisks indicate significant differences in mutant expression compared to the respective WT, as determined by least-squares means Dunnett’s test. Means correspond to five biological replicates, and the error bars show SEM. Statistical significance for each sample was represented in the figures. Dunnett’s test results were displayed as asterisks (*p < 0.05, **p < 0.01, ***p < 0.001). Black stars represent differences in expression in infected mutants compared to infected WT, and red stars indicate significant differences in expression between non-infected and corresponding infected samples.

**Figure 6:**
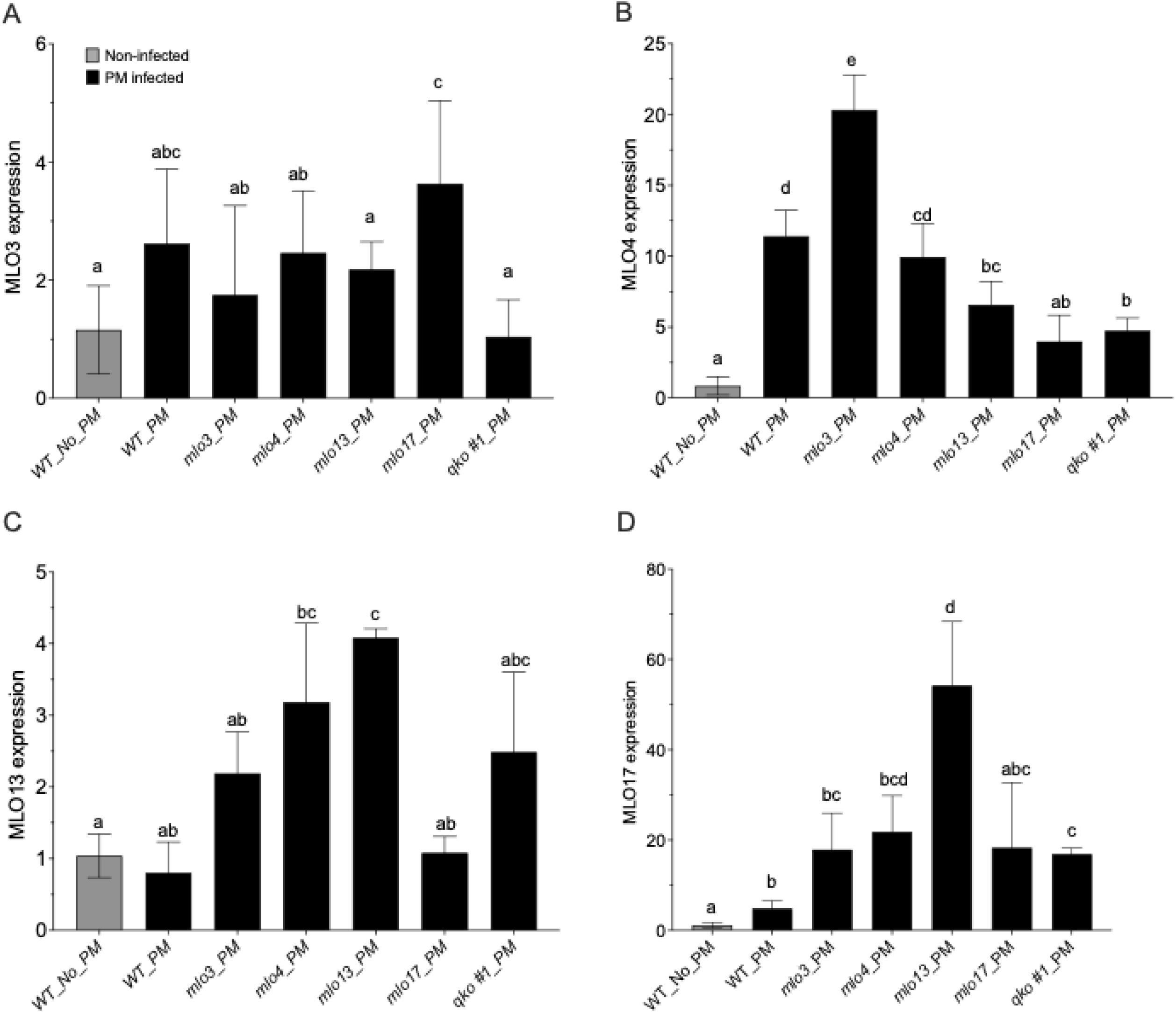
Late transcriptional response of *MLO* orthologs in WT and *mlo* mutants 2wpi through quantitative RT-PCR analyses. The results are expressed as fold changes relative to expression in non-infected WT. **A,B,C,D**. Fold-Change expressions relative to WT infected with *Erysiphe necator* in *MLO3*, *4*, 1*3*, and *17*, respectively. To assess statistical significance, model assumptions were evaluated using residual diagnostics. While most genes showed no substantial deviations, mild heteroscedasticity was detected for *MLO13*; therefore, inference was based on heteroscedasticity-robust standard error. Post-hoc Tukey’s HSD analysis was performed to assess the expression differences between infected mutants and WT control, as well as WT-infected leaves. Levels not connected by the same letters are significantly different.

The long-term expression response of *MLO3* at 2 wpi was not different from that of infected WT in any mutant line (**Fig. 6A**). *MLO4* expression was maintained at a higher level in infected WT compared to non-infected plants (∼10FC), and so was in infected *mlo3*, *mlo3,* and *mlo13*. However, *MLO4* expression was maintained at a much higher level in *mlo3* compared to WT (**Fig. 6B**). While the early-stage induction of *MLO13* in *mlo3* and *mlo13* was maintained at 2 wpi stage, it was also expressed at a higher level in PM-challenged *mlo4* at this stage (**Fig. 6C**). The *MLO17* gene is by far the most highly induced in the mutants during the last stages of disease (**Fig. 6D**). Consistent with its close genetic proximity with *MLO13*, significantly higher expression was observed in infected *mlo13* with 50-fold higher level (**Fig. 6D**). Taken together, these results indicate that *MLO4* might be expressed at higher rate in response to the loss of MLOs 3 and 13 during the early infection, and the high expression level is maintained until the mature stages of infection in *mlo3*. By contrast, *MLO13* is the most responsive gene to mutations in the other homologs at either the early or late stages of infection. MLO17 expression increased specifically in the chimeric mlo13 line due to the partial loss of its paralog.

## Discussion

The induction of *VvMLOs 3*, *4*, *13*, and *17*, which represent the closest orthologs of *AtMLOs 2*, *6*, and *12*, followed previously reported trends in grapevine during PM infection (Feechan et al., 2008; Winterhagen et al., 2008). The early and late expression peaks indicate potential roles for MLOs 4 and 17 during initial fungal establishment, and for MLOs 4, 13, and 17 in later stages of fungal growth. Relative transcript abundance of *MLOs 13* and *17* was more than 50-fold higher compared to *MLOs 3* and *4* at 72 hpi, indicating their major role in PM-susceptibility (**Fig. 1B**). Tissue-specific expression studies reported *MLOs 3* and *4* are highly expressed in grapevine floral tissues, suggesting their likely involvement in the flowering process, while MLOs 13 and 17 have leaf-specific expression (Feechan et al., 2008; Winterhagen et al., 2008). Altogether, the expression peaks of the four MLOs spanned over the entire PM pathogenesis process (up to 2wpi), and mutations in these MLOs effected the success of key PM-infection stages, such as sporulation and hyphal growth.

The multiplexed editing strategy described in this report, which targeted multiple MLOs, heavily depended on relative sgRNA efficiencies *in vivo* (**Fig. S5**). This was evidenced by the fact that only two *qko* lines had all four *MLOs* edited out of 32 potential qko transformants genotyped. As a result, incidental *mlo* mutant combinations could be retrieved primarily from the qko mutant pool. In addition, most genotyped mutants exhibited a complex or mosaic editing pattern comprising several differentially edited cell lineages, as well as chimeric mutants with WT alleles (**Table 1**). Mosaic editing and WT chimerism might originate from various events, including the CRISPR plasmids being introduced at or after the single-cell stage, a delayed Cas9 activity, and a random editing of cells at different stages in the proliferating embryogenic structure (Cui et al., 2021; Impens et al., 2022; Mehravar et al., 2019). Nevertheless, the differential editing rate of one or more MLOs enabled us to understand the roles of different *mlo* combinations in PM susceptibility, *MLO*-resistance, and *MLO*-associated pleiotropic effects.

The effect of having a proportion of functional MLO alleles coexisting in WT chimeric mutants is seen in *mlo13* and *mlo17,* having 25% target-editing rate (mutants composed of 25% cells with mutation and 75% WT functional allele), which supported significant hypervirulent growth of pathogens (**Fig. 3**). We hypothesize that the cell-autonomous nature of mlo resistance and the interspersed mix of edited and unedited cells in the leaf might have led to hyphal foraging into wider areas increasing the mycelial spread but not sporulation. It is also possible that partial loss of MLO function results in higher expression of close paralog, making the plant more susceptible. Contrary to partial editing, complete editing of *MLO17,* together with *MLO13,* in *qko#1* led to restricted mycelial growth, implicating MLOs 13 and 17 in promoting mycelial growth. (**Fig. 2 and 3**). Pathogenicity assays in *mlo* mutant combinations also established a combined role of MLOs 3 and 4 in fungal sporulation as *mlo3,4*, and other lines containing this combination of edited *MLOs* showed significantly reduced sporulation (**Fig. 3**). While MLOs were thought to be mainly involved in the process of host cell penetration and fungal establishment, reduced hyphal growth or conidiophore production caused by *mlo* mutations in *Arabidopsis* and tomato were reported, indicating a post-penetration role for MLOs during fungal development (Acevedo-Garcia et al., 2017; Consonni et al., 2006b).

Previous MLO studies in grapevine that generated non-chimeric, fully edited lines through gene editing reported partial to significant reductions in PM disease severity with single and double mutations in MLO3 and MLO13 (Dong-Yan et al., 2020; Moffa et al., 2024). In contrast, the single *mlo* mutants generated in our study showed no PM resistance, including the fully edited and non-chimeric *mlo3* and *mlo4* (**Fig. 4**). Likewise, *mlo13,17*, in which MLO17 was edited incompletely, showed no significant PM resistance. Fully edited *mlo13,17* line from our experiments exhibited severe pleiotropic effects, including black root, loss of root gravitropism, and narrow leaves, and did not survive. Other double KO lines showed partial PM resistance, as evidenced by reduced fungal biomass in infected leaves, either due to inhibited hyphal growth or sporulation (**Fig. 4 and 5**). Mutations in all four MLOs resulted in high PM resistance, but *qko#1* line exhibited complete resistance while *qko#2* line showed partial resistance. Complete PM-resistance in *qko#1* was a result of complex editing in which all the cells in the mutant have edited *MLOs 3* and *13*, 12% cells contain unedited WT *MLO 17* allele, and 80% cells with WT *MLO4* allele, and the editing in all four *MLOs* was of a mosaic nature (**Fig. 2 and S4**). Besides being completely PM-resistant, the qko#1 line was also devoid of any *mlo*-associated pleiotropic defects, often seen in null mutations. *qko#2* line differs from *qko#1* in having MLO3 and 4 with complete editing and low levels of editing in MLOs 13 and 17 due to WT chimerism (**Fig. 2**). Comparison of *qko* genotypes with different editing patterns indicates greater contribution of *mlo17* allele to resistance, followed by *mlo 3* and *13* alleles, consistent with the unequal genetic redundancy of MLOs reported in *Arabidopsis* with respect to PM resistance and developmental defects (Consonni et al., 2006b, 2010). Similar *mlo*-based resistance with alleviated pleiotropic effects was also reported in wheat due to a weaker mutation of a homolog (Ingvardsen et al., 2019). In addition, PM resistance with reduced pleiotropic effects was associated with mosaic editing, together with WT chimerism, in grapevine *edr1* mutants, whereas pure homozygous and bi-allelic *edr1* lines suffer from developmental defects (Yu et al., 2024).

If the aim is to generate grapevine PM-resistant lines through null mutations in MLOs, it may come with high agronomic costs because we found a certain level of chimerism, and partial MLO activity is an essential trade-off to ensure survival of the PM-resistant mutant lines that are knocked out for susceptibility genes (Yu et al., 2024). Most MLO mutants in our study showed growth and developmental abnormalities, and the complex nature of the editing (partial, mosaic, and chimeric events), along with the likely functional overlap between MLO homologs have made it challenging to accurately attribute the observed pleiotropic effects to any single MLO or combinations (**Table 1**). It is possible that different levels of WT chimerism observed in the mutant lines contribute to WT functional MLO alleles, which partly reduce the developmental defects. Further, pleiotropic phenotypes associated with *mlo* mutants are also known to be heavily influenced by genotype-by-environment interaction (Freh et al., 2024) (Brown & Rant, 2013; McGrann et al., 2014). Common pleiotropic effects include spontaneous necrotic lesions, leaf senescence, and stunted growth, which we observed (**Table 1**), are consistent with reported roles of MLOs in cell death protection and in delaying leaf senescence, in addition to their function as negative regulators of powdery mildew resistance (Piffanelli et al., 2002). Only three mutant lines, *mlo3*, *mlo3,13,* and *qko#1,* lacked pleiotropic effects caused by the loss of MLO genes and showed similar growth, flowering, and fruit development as WT plants. Observation of developmental defects versus PM resistance in genotypes derived from gene-editing events supports the assertion that a certain degree of chimerism in the mutants might be beneficial for survival when targeting genes with essential functions in other developmental processes (Mehravar et al., 2019)(Piffanelli et al., 2004). Likewise, RNAi knockdown of all four MLOs, with 50% reduced expression, did not result in any pleiotropic effects in grapevine (Pessina, Lenzi, et al., 2016b). In our study, Generally, the mutants with non-edited or incompletely edited *MLO4*, were free of developmental defects, including *qko#1* (Fig. 2 and **Table 1**) suggesting a predominant role of MLO4 in vegetative and reproductive development beyond its accessory role in PM-susceptibility. After three cycles of growth under greenhouse conditions, all these mutants free of developmental effects show no differences in growth, flowering, and fruit production compared with WT plants, suggesting that either a long-term compensation mechanism has been established or that partial editing of *MLO4* ensures minimal production of the native MLO4 protein, thereby preventing any major long-term growth defect while slightly improving the resistance to PM (**Fig. 3 and Fig.4**).

Our study also supports functional redundancy in clade-V MLOs, which could also explain both the lack of PM-resistance observed in mutants and the absence of pleiotropic effects following edit(s) on one or multiple MLO genes. The compensatory influence of one or the other homologs might partially rescue the PM susceptibility phenotype or alleviate pleiotropic effects when MLOs, which share overlapping developmental functions, are lost. Besides, functional redundancy among MLOs can span across phylogenetic clades. *At*MLO2 (clade V) and *At*MLO7 (clade III) can each partially compensate for the loss of the other’s function in PM susceptibility and pollen tube growth, respectively (Li & Xiao, 2025). At the molecular level, unequal redundancy often results from active compensation mechanisms between homologs (Iohannes & Jackson, 2023). In the context of PM-susceptibility, our data suggest a compensatory influence of MLO4 for the loss of MLO3, and of MLO13 for the loss of MLO3 (**Fig. 6 B, C**). Both MLO13 and MLO17 appear to exhibit a long-term compensatory influence for the loss of other homologs (**Fig. 7 C, D**). Screening the other MLO-related genes in these mutants, challenged by PM, could reveal the involvement of other MLO genes outside clade V.

Altogether, mutations in grapevine MLOs 3, 4, 13, and 17 contribute unequally to PM resistance, and a balanced mutant genotype containing one or more partially functional homologs that control other plant developmental processes is required to achieve PM resistance without severe detrimental effects. Comparing the complete and partial resistance exhibited in *qko* lines *#1* and *#2,* and their editing signatures, indicates that the contribution of the MLO17 gene to susceptibility is greater than the contribution of the *MLO3* and *MLO13* genes. This study also revealed additional roles for *MLO4*, *13*, and *17* beyond PM susceptibility. The inability to maintain the *mlo13,17* line under tissue culture conditions might support the notion that *MLO13* and *17* are paralogs and co-involved in root development.

---

**Figure S1:** Schematic illustration of the Single Transcript Unit transgenic cassette used for the expression of the CRISPR-Cas9 gene with the sgRNA array.

**Figure S2:** Specificity of sgRNAs across the *MLO* gene targets. Each sgRNA was tested for cleavage activity on all four MLOs in an *in vitro* cleavage assay. Respective full-length *MLO* genes cloned into the pGMT vector were used as template DNA along with specific sgRNAs and Cas9 protein in the *in vitro* cleavage reaction. Target *MLO* gene-containing plasmids are linearized (∼5kb), while the non-target MLO plasmids show nicked relaxed (slower migration), and supercoiled forms (faster migration) of the circular plasmids.

**Figure S3.** Stereoscopic observation of germination and growth of *E. necator* spores on infected leaves of WT microvine leaves and of *mlo* mutants 24hpi.

**Figure S4.** Comparative analysis of the editing performance deduced from DECODR analysis and long-read sequence technology.

**Figure S5.** Genotyping Analysis of the four *mlo* genes on qko#1 mutant using long-read sequencing. Alignment on the target sequence (sgRNA sequence + PAM). For each sequence identified, the percentage represents the number of reads found relative to the total number of reads, accounting for sequencing errors.

**Table S1**. Single guide RNA sequences

**Table S2**. PCR and sequencing primers used for the genotyping of MLO mutants.

**Table S3**. *MLO* gene nomenclature used in this study and cross-referenced with alternative names used in other studies. *^1^* Feechan et al. (2008) - Followed in the current study; *^2^* Winterhagen et al. (2008) and Pessina et al. (2016); *^3^* Canaguier et al., 2017; *^5^* http://urgi.versailles.inra.fr/Species/Vitis/Data-Sequences/Genome-sequences.

**Table S4**. Timeline of the critical infection stages of *Erisyphe necator* in grapevine leaves.

**Table S5**. Overall efficiency of editing for the four *MLO* genes calculated for each *mlo* gene in the mutants.

**Table S6**. Overall performance of CRISPR-mediated editing based on the number of edited transgenic lines.

