## Supplemental for "CRISPR-Cas9 editing of grapevine clade V-related *mlo* genes revealed new insights into the contribution of each *mlo* gene to grapevine powdery mildew resistance (*Erysiphe necator*) and plant development"

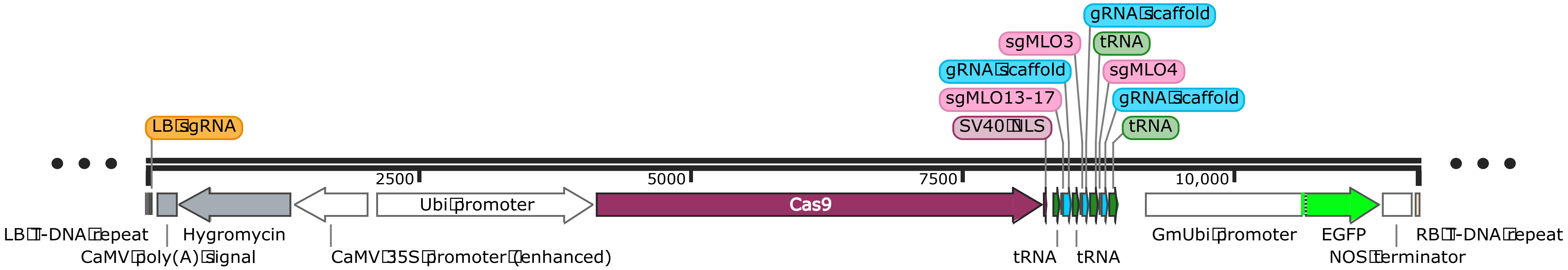
gRNA scaffold
sgMLO3
tRNA
gRNA scaffold
sgMLO4
sgMLO13-17
gRNA scaffold
SV40 NLS
tRNA
LB sgRNA
2500
5000
7500
10,000
Ubi promoter
Cas9
LB T-DNA repeat
Hygromycin
CaMV poly(A) signal
CaMV 35S promoter (enhanced)
tRNA
tRNA
GmUbi promoter
EGFP
NOS terminator
RB T-DNA repeat

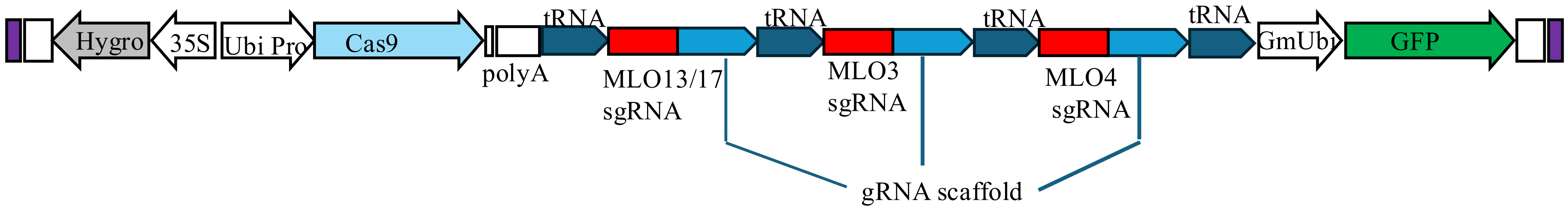
Hygro
35S
Ubi Pro
Cas9
polyA
tRNA
tRNA
tRNA
tRNA
MLO13/17
sgRNA
MLO3
sgRNA
MLO4
sgRNA
gRNA scaffold
GmUbi
GFP

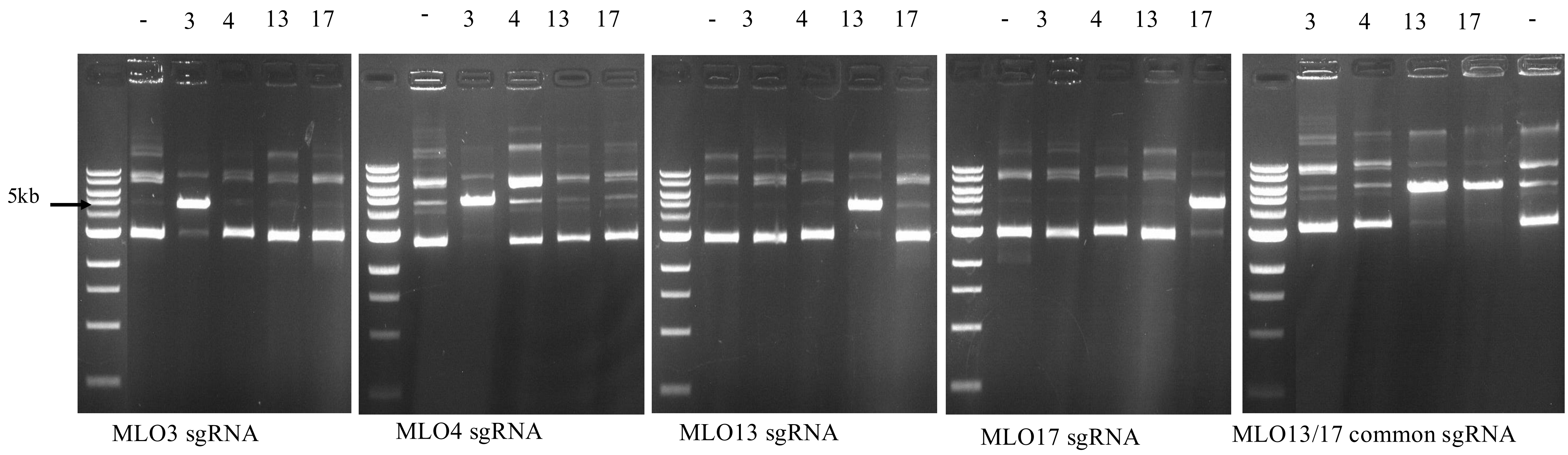
- 3 4 13 17
- 3 4 13 17
- 3 4 13 17
- 3 4 13 17
3 4 13 17 -
5kb
MLO3 sgRNA
MLO4 sgRNA
MLO13 sgRNA
MLO17 sgRNA
MLO13/17 common sgRNA

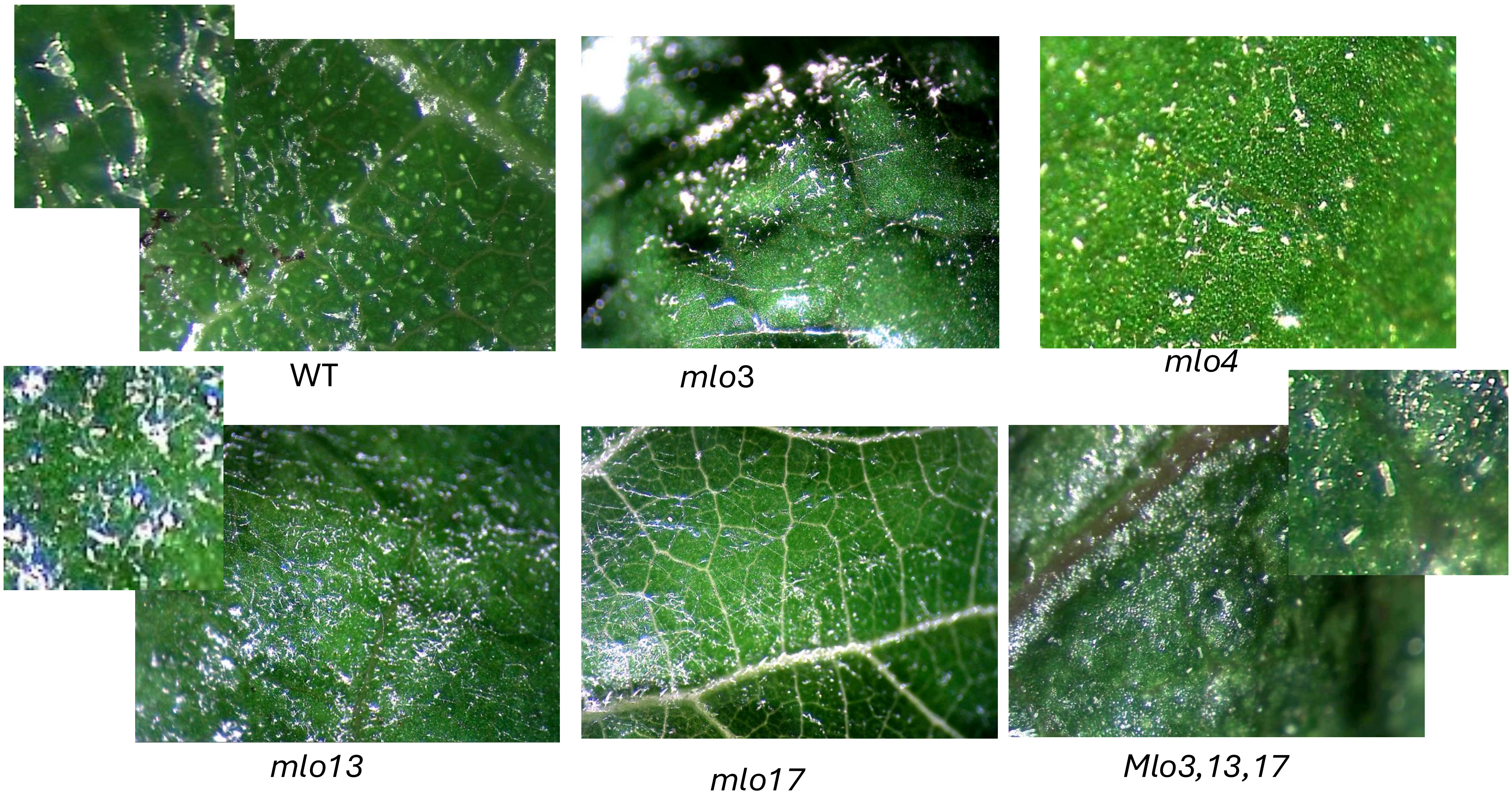
WT
mlo3
mlo4
mlo13
mlo17
Mlo3,13,17

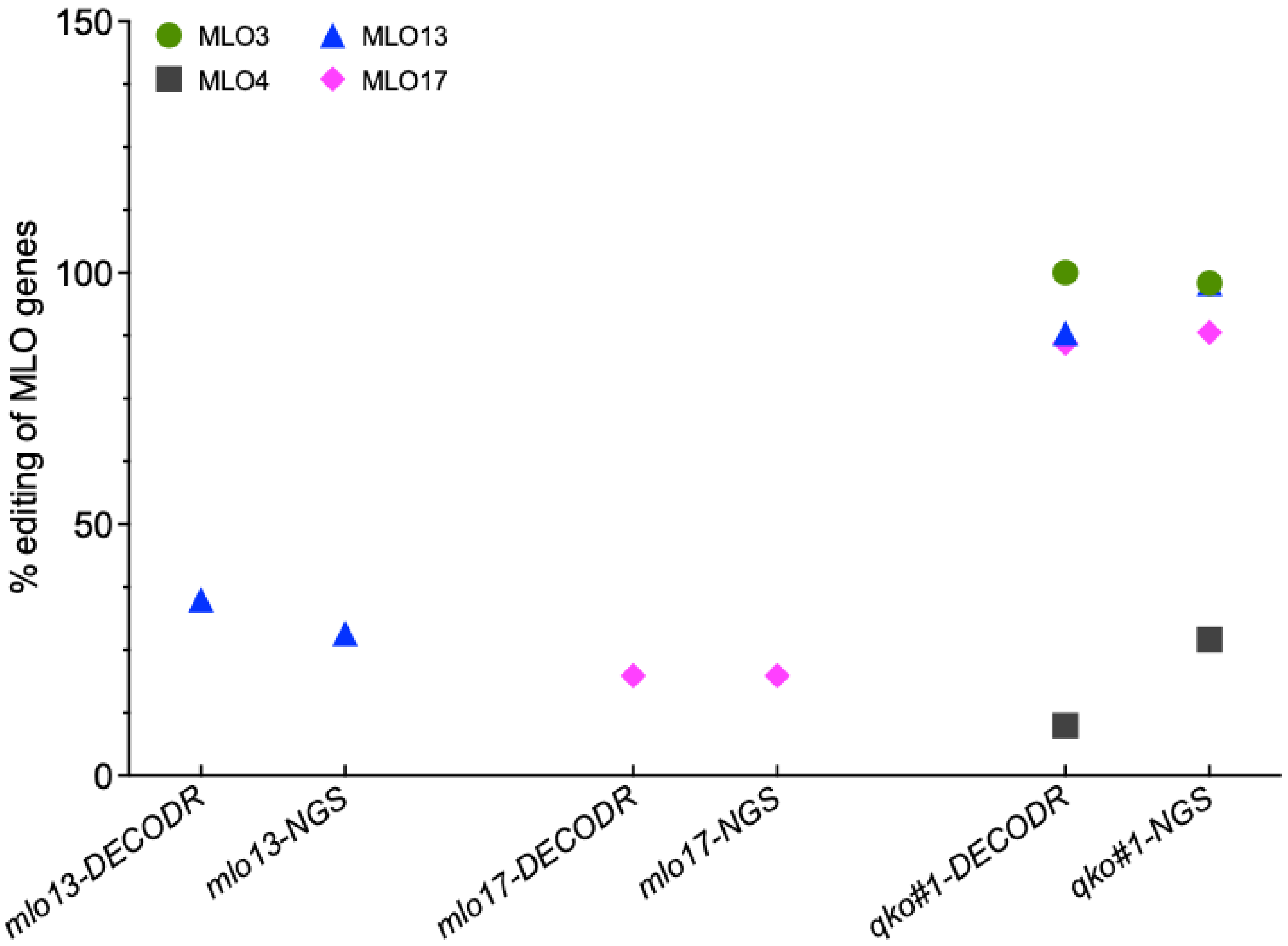
150
100
50
0
% editing of MLO genes
MLO3
MLO4
MLO13
MLO17
mlo13-DECODR
mlo13-NGS
mlo17-DECODR
mlo17-NGS
qko#1-DECODR
qko#1-NGS

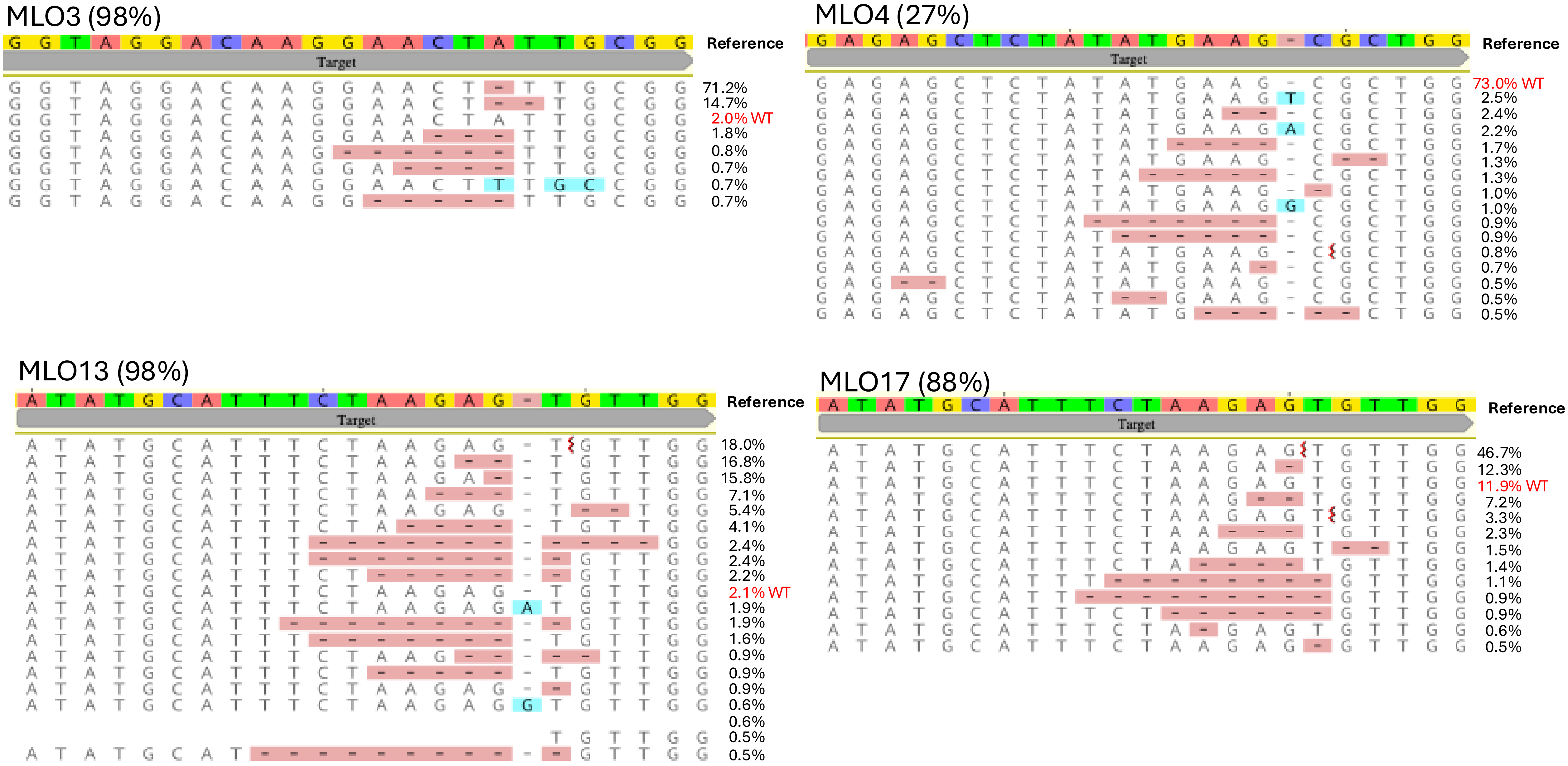
MLO3 (98%)
GGTAGGACAAGGAACTATTGCGG
Reference
Target
71.2%
14.7%
2.0% WT
1.8%
0.8%
0.7%
0.7%
0.7%
MLO4 (27%)
GAGAGCTCTATATGAAG-CGCTGG
Reference
Target
73.0% WT
2.5%
2.4%
2.2%
1.7%
1.3%
1.3%
1.0%
1.0%
0.9%
0.9%
0.8%
0.7%
0.5%
0.5%
0.5%
MLO13 (98%)
ATATGCATTTCTAAGAG-TGTTGG
Reference
Target
18.0%
16.8%
15.8%
7.1%
5.4%
4.1%
2.4%
2.4%
2.2%
2.1% WT
1.9%
1.9%
1.6%
0.9%
0.9%
0.9%
0.6%
0.6%
0.5%
0.5%
MLO17 (88%)
ATATGCATTTCTAAGAGTGTTGG
Reference
Target
46.7%
12.3%
11.9% WT
7.2%
3.3%
2.3%
1.5%
1.4%
1.1%
0.9%
0.9%
0.6%
0.5%

| Target Gene | sgRNA | PAM | Strand | Position from ATG | Exon | CRISTA score *(Abadi et al. 2017)* |
|---|---|---|---|---|---|---|
| *MLO3* | GGTAGGACAAGGAACTATTG | CGG | + | 411-430 | 3rd | *0.869* |
| *MLO4* | GAGAGCTCTATATGAAGCGC | TGG | + | 239-258 | 2nd | *0.934* |
| *MLO13* | GAAATGCATATCTTTGAAAT | GGG | - | 815-795 | 3rd | *0.900* |
| *MLO17* | CACTTGGCACCCTTGTAAAA | AGG | + | 1114-1133 | 3rd | *0.903* |
| Common *MLO13/17* | ATATGCATTTCTAAGAGTGT | TGG | + | 805-824 | 3rd | *0.907* |

Genotyping primers

| **Mutant** | **Forward PCR primer (5'-3')** | **Reverse PCR primer (5'-3')** | **Nested Sequencing primer** |
|---|---|---|---|
| *MLO3* | TAAGGGATCAAAGGA TCGA | CCTTCTCTTGCCATAATCT | ACGAGCTCTTTATGAAGCAC |
| *MLO4* | GGCAACCGGAGGAAGGTCG | TCTCTTCTTCAGACTTGCTGC | GTACATCATTCATCTCACTG |
| *MLO13* | TGGTTAGCAAGAAGAAACAA | GAGCTGATGGATCCCATAG | TGGCTTCTTCTCATTGAGA |
| *MLO17* | TGGTTAAAAGGCAGACACAGG | GAGTTGATCAATCCCATAGG | TCATTGAGAGTAGTTCTTCC |

Real-time PCR primers

| Gene | Forward primer (5'-3') | Reverse primer (5'-3') |
|---|---|---|
| MLO3 | CAGCCCATGGACTTTTCATTCG | TTCGTTGCCCACAGCTATAC |
| MLO4 | AGCCAACGATCTTCCATGAG | GGCCATCTTGACATGGGTGT |
| MLO13 | GAAAGCCTCTCTCTCTCTATTTCC | TTTCTGCTGGTGCTCAGCTCTC |
| MLO17 | AGGGACAAGGGATTGGAGAC | AAATACAAGCCGCAGAAAGG |
| VvActin1 gene | GTGTGGAGGGATTTATCTGTAATG | CAATCACTCTCCTGCTACAAAC |
| *E. necator* UNC gene | CCGCCAGAGACCTCATCCAA | TGTGCGTTCAAACATTCGATGA |

| Gene name[1] | Alternative name[2] | V3-COST-annotation[3] |
|---|---|---|
| *VvMLO3* | *VvMLO11* | Vitvi08g01055 |
| *VvMLO4* | *VvMLO13* | Vitvi06g00308 |
| *VvMLO13* | *VvMLO6* | Vitvi13g00579 |
| *VvMLO17* | *VvMLO7* | Vitvi13g00578 |

| Stages | Earliest Time Post-Inoculation (hours or days) | Approximate TImelength | References |
|---|---|---|---|
| Conidial Adhesion | Minutes following the contact | Conidia rapidly adhere on the leaf surface through hydrophobic interactions and extracellular adhesive material | Gadoury et al., 2012 |
| Conidial Germination | 2 hpi | Germination begins within hours under favorable humidity/ temp | Gadoury et al., 2012 |
| Appressorium & Penetration | 6 hpi | Histological observations by 24-48 hpi have appressoria/haustoria visible | Feechan et al., 2010 |
| Haustorium formation | 24 hpi | Gene expression atlas identifies haustorial structures by 24 hpi | Bonarota et al., 2025 |
| Hyphal Growth & colony expansion | 48 hpi | Progressive epiphytic mycelium visible by 2-3 dpi in controlled inoculations | Gadoury et al., 2012 |
| Sporulation/latent period | 5 dpi | New conidia produced 5-10 dpi under optimal conditions | Gadoury et al., 2012 |

| **Mutant** | **Gene** | **Indel** | **Editing %** | **Frame shift mutation %** | ***$R^2$*** |
|---|---|---|---|---|---|
| mlo3 | *MLO3* | -1 | 100 | 100 | *1.00* |
| mlo3 | *MLO4* | +1 | 80 | 80 | *0.99* |
| mlo13 | *MLO13* | +1, -3, -84 | 35, 9, 8 | 35 | *0.62* |
| mlo17 | *MLO17* | -1, -4, -3 | 15, 10, 8 | 25 | *0.97* |
| mlo3,4 | *MLO3*<br>*MLO4* | -2<br>+1 | 100<br>41 | 100<br>41 | *1.00*<br>*0.98* |
| mlo3,13 | *MLO3*<br>*MLO13* | -1<br>+1, -1, -3 | 100<br>36, 11, 9 | 100<br>47 | *0.06*<br>*0.70* |
| mlo13,17 | *MLO13*<br>*MLO17* | +1, -1, -3<br>-2 | 36, 12, 8<br>8 | 48<br>8 | *0.72*<br>*0.95* |
| mlo3,13,17 | *MLO3*<br>*MLO13*<br>*MLO17* | -1, -2<br>-1, -2, +1<br>+1, -1, -2 | 58, 31<br>38, 27, 26<br>12, 14, 22 | 89<br>92<br>48 | *0.98*<br>*0.93*<br>*0.96* |
| mlo3,4,13,17 # 1 | *MLO3*<br>*MLO4*<br>*MLO13*<br>*MLO17* | -1<br>+2 (0)<br>+2, +1, -1<br>-1, -2 | 94<br>51 (100) **(20)**<br>16, 13, 12<br>18, 8 | 94<br>51 (0)<br>41<br>26 | *0.97*<br>*0.98 (1.00)*<br>*0.97*<br>*0.98* |
| mlo3,4,13,17 #2 | *MLO3*<br>*MLO4*<br>*MLO13*<br>*MLO17* | -1, -2<br>+1<br>+1, -1, -2, -3<br>+1, -1, -2 | 18, 82<br>10 **(30)**<br>30, 26, 32, 12<br>63, 17, 9 | 100<br>10 **(30)**<br>88<br>89 | *0.98*<br>*0.99*<br>*0.83*<br>*0.98* |

| Gene | Number of transgenic lines screened | Number of transgenic lines with target edited | Efficiency % |
|---|---|---|---|
| *MLO3* | 61 | 19 | 31 |
| *MLO4* | 65 | 8 | 12 |
| *MLO13* | 57 | 13 | 22 |
| *MLO17* | 64 | 11 | 17 |
